# Inducing a meditative state by artificial perturbations: A causal mechanistic understanding of brain dynamics underlying meditation

**DOI:** 10.1101/2023.07.27.550828

**Authors:** Paulina Clara Dagnino, Javier A. Galadí, Estela Càmara, Gustavo Deco, Anira Escrichs

## Abstract

Contemplative neuroscience has increasingly explored meditation using neuroimaging. However, the brain mechanisms underlying meditation remain elusive. Here, we implemented a causal mechanistic framework to explore the spatiotemporal dynamics of expert meditators during meditation and rest. We first applied a model-free approach by defining a probabilistic metastable substate (PMS) space for each state, consisting of different probabilities of occurrence from a repertoire of dynamic patterns. Different brain signatures were mainly found in the triple-network model (i.e., the executive control, salience, and default-mode networks). Moreover, we implemented a model-based approach by adjusting the PMS of the resting state to a whole-brain model, which enabled us to explore *in silico* perturbations to transition to the meditation state. Consequently, we assessed the sensitivity of different brain areas regarding their perturbability and their mechanistic local-global effects. Using a synchronous protocol, we successfully transitioned from the resting state to the meditative state by shifting areas mainly from the somatomotor and dorsal attention networks. Overall, our work reveals distinct whole-brain dynamics in meditation compared to rest, and how the meditation state can be induced with localized artificial perturbations. It motivates future work regarding meditation as a practice in health and as a potential therapy for brain disorders.

## Introduction

> If we transform our way of perceiving things, then we transform the quality of our lives. This kind of transformation is brought about by the form of mind training known as meditation.

Matthieu Ricard (Ricard and Wolf, 2017)

Meditation encompasses a wide range of mental training techniques that allow to explore one’s inner self and relation with the outer world (Kabat-Zinn, 2005; Hanh, 1991). It has roots in different cultures and religions, with its origin dating back to ancient Hindu and Buddhist spiritual traditions in India (Sharma, 2015; Wynne, 2009; Dumoulin, 1988). The different practices of meditation (e.g., attentional, open monitoring, loving kindness) have a variety of positive effects, including present-moment awareness and observation of experience with openness, acceptance, and non-attachment (Dahl et al., 2015; Ricard et al., 2014; Lutz et al., 2009; Kabat-Zinn, 2003). They can also promote stress reduction, well-being, social connectedness, self-awareness, and attention and emotional regulation (Basso et al., 2019; Champion et al., 2018; Pascoe et al., 2017; Guendelman et al., 2017; Tang et al., 2015; Marchand, 2012; Ludwig and Kabat-Zinn, 2008; Valentine and Sweet, 2007).

Over the past years, meditation is increasingly being explored as an intervention to enhance well-being and alleviate suffering in the healthy general population, as well as to prevent and recover from disability and disease (Dahl et al., 2015; Ludwig and Kabat-Zinn, 2008). The field of contemplative neuroscience, in particular, uses neuroscience tools – e.g., functional magnetic resonance imaging (fMRI) – to study the effects of meditation (Davidson R, 2013; Varela, 1993). Neuroanatomical studies have investigated structural gray matter thickness in meditators relative to controls, finding structural evidence such as the thickening of the prefrontal cortex with experience in long-term meditation practice (Lazar et al., 2005). Moreover, functional brain imaging studies have largely shown that key brain areas relevant to meditation belong to the default mode network (DMN), the salience network and the control network (Ganesan et al., 2022; Sezer et al., 2022; Fox et al., 2016; Hasenkamp et al., 2012). Interestingly, these resting-state networks belong to the triple-network model hypothesis (Menon, 2011). Different static and seed-based approaches have analyzed blood oxygen level-dependent (BOLD) signal activity and network connectivity changes of meditation in health (Vishnubhotla et al., 2021; Martínez et al., 2021; Kyeong et al., 2017; Garrison et al., 2014; Ives-Deliperi et al., 2013; Pagnoni, 2012; Kilpatrick et al., 2011) and brain disorders (Li et al., 2022; Datko et al., 2022; Lifshitz et al., 2019; Goldin et al., 2009). Furthermore, theoretical methods have considered the underlying brain dynamics (Mooneyham et al., 2017; Bremer et al., 2022; Martínez et al., 2021; Lim et al., 2018; Marusak et al., 2018), including whole-brain approaches such as turbulent dynamics (Escrichs et al., 2022), dynamical complexity (Escrichs et al., 2019) and structural and effective connectivity (De Filippi et al., 2022). Whole-brain dynamics analysis and computational modeling can shed light on the mechanisms underlying meditation practices (Tang et al., 2015). Nonetheless, they remain largely unexplored.

The elegant awakening framework proposed by Deco et al. (2019) has been robust and helpful in elucidating the underlying causal mechanisms and brain transitions in different brain states towards control regimes (Escrichs et al., 2023; Vohryzek et al., 2022; Mana et al., 2023; Deco et al., 2019). It consists of a model-free approach, Leading Eigenvector Dynamics Analysis (LEiDA) (Cabral et al., 2017), and a model-based approach, consisting of Hopf whole-brain computational models and off-line *in silico* perturbations. Brain dynamics are studied with reference to the concept of metastability, which corresponds to the ability of a system to maintain equilibrium regardless of slight perturbations (Freyer et al., 2012; Kelso, 2012). LEiDA framework, in particular, typifies a brain state with its probabilistic metastable substate (PMS) space (Escrichs et al., 2021; Kringelbach and Deco, 2020; Figueroa et al., 2019; Lord et al., 2019; Deco et al., 2019; Cabral et al., 2017) Each PMS corresponds to a discrete repertoire of metastable substates (i.e., dynamic patterns) at critical points between chaos and order (Cabral et al., 2017; Deco et al., 2017b), in which substate duration and arrangement is a dynamic signature of a particular brain state (e.g., sleep, anesthesia) (Deco and Kringelbach, 2016; Tognoli and Kelso, 2014). By building a whole-brain model composed of a network of coupled local nodes (Botvinik-Nezer et al., 2020; Deco et al., 2019), the empirical PMS can be simulated which then allows to force a transition to a desired control state with *in silico* external stimulation. Hence, whole-brain reactivity and its causal underpinnings can be studied by stimulating one brain area at a time (Kringelbach and Deco, 2020; Breakspear, 2017; Deco et al., 2015; Deco and Kringelbach, 2014).

Using the framework outlined above, we investigated the underlying causal mechanisms of brain activity of expert meditators during a meditation focused on breath (i.e., Anapanasati in Pali language) and resting-state. As a first step, we applied LEiDA to define the PMS of meditation and rest. As a second step, we developed a Hopf whole-brain model based on the empirical PMS of resting-state at the bifurcation point, where oscillatory and noisy regimes are undifferentiated, and the system is critical. Lastly, we applied off-line *in silico* external unilateral and localized probing to force the transition from the resting-state PMS to the meditation PMS. Thus, we could evaluate local brain areas in terms of their sensitivity to stimulation and their mechanistic global effects after perturbation. Overall, our work contributes to the state-of-the-art of contemplative neuroscience, as we try to understand brain mechanisms during meditation and resting-state in expert meditators. Furthermore, it motivates current and potential future therapies based on meditation (Ludwig and Kabat-Zinn, 2008; Grossman et al., 2004). This way, benefit actual and future practices for the healthy population and brain disorders.

## Materials and Methods

### Participants

This study selected 20 experienced meditators and 20 healthy controls from a dataset previously described in Escrichs et al. (2019). The meditator group consisted of 13 males and 7 females (mean age=39.8 years (SD=10.29); education=13.6 years; and meditation experience=9,526.9 hours (SD=8,619.8). Meditators were recruited from Vipassana communities in Barcelona, Catalonia, Spain, and had more than 1,000 hours of meditation experience with a daily practice of over one hour. The healthy control group consisted of 13 males and 7 females with no prior meditation experience (mean age= 39.75 years (SD=10.13); education=13.8 years). Both groups were well-matched for age, gender, and educational level and reported no history of neurological disorder. All participants provided written informed consent, and the study was approved by the Ethics Committee of the Bellvitge University Hospital in accordance with the Helsinki Declaration.

### MRI Data Acquisition

Magnetic resonance imaging (MRI) scans were performed on a 3T (Siemens TRIO) scanner using a 32-channel receiver coil. High-resolution T1-weighted images were acquired with 208 contiguous sagittal slices, with the following parameters: repetition time (TR) 1970 ms, echo time (TE) 2.34 ms, inversion time (IT) 1050 ms, flip angle 9°, field of view (FOV) 256 mm, and isotropic voxel size 1 mm. Resting-state and meditation functional MRI (fMRI) scans were obtained using a single-shot gradient-echo echo-planar imaging (EPI) sequence, comprising a total of 450 volumes, with TR 2000 ms, TE 29 ms, FOV 240 mm, in-plane resolution 3 mm, 32 transversal slices, thickness 4 mm, and a flip angle of 80°. Diffusion-weighted imaging (DWI) data were collected using a dual spin-echo diffusion tensor imaging (DTI) sequence, comprising 60 contiguous axial slices, with TE 92 ms, FOV 236 mm, isotropic voxel size 2 mm, no gap, and matrix sizes 118 x 118. Diffusion was measured with 64 optimal non-collinear diffusion directions using a single b value of 1,500 s/mm2 interleaved with 9 non-diffusion b0 images, with a frequency-selective fat saturation pulse applied to avoid artifacts.

### Experimental conditions

During rest, participants were instructed not to think about anything in particular. During meditation, participants had to focus on their natural breathing and come back to it whenever the mind wandered. Participants were asked to remain still and move as little as possible inside the scanner. Consistent with previous methodologies in meditation neuroimaging studies (Braboszcz et al., 2017; Ives-Deliperi et al., 2011; Manna et al., 2010; Davanger et al., 2010), we excluded the first and last volumes of the fMRI conditions in the final contrast analyses to ensure that the initial and final stages of the acquisition did not influence the physiological measures collected during the meditation and resting-state periods. These excluded periods, comprising the first and last 110 timepoints, served as a build-up period for the meditation practice, as the effects of meditation have been observed to accumulate slowly over time. This approach was taken to avoid potential confounding effects that may arise from the initial and final stages of the meditation practice.

### Resting-state fMRI preprocessing

Preprocessing was done using MELODIC version 3.14 (Beckmann and Smith, 2004), which is part of FMRIB’s Software Library (FSL, http://fsl.fmrib.ox.ac.uk/fsl). First, the initial 5 volumes were discarded, followed by motion correction using MCFLIRT (Jenkinson et al., 2002), non-brain removal using BET (Brain Extraction Tool) (Smith, 2002), spatial smoothing with 5 mm FWHM Gaussian Kernel, rigid-body registration, high pass filter cutoff = 100.0s, and single-session ICA with automatic dimensionality estimation. Noise removal was performed using FIX (FMRIB’s ICA-based X-noiseifier) (Griffanti et al., 2014), which independently removes noise components for each participant. FSLeyes in Melodic mode was used to manually classify the single-subject Independent Components (ICs) into “good” for signal, “bad” for noise, and “unknown” for ambiguous components, based on the spatial map, the time series, and the temporal power spectrum (Griffanti et al., 2017; Salimi-Khorshidi et al., 2014). Finally, FIX was applied using the default parameters to obtain a cleaned version of the functional data.

We then used FSL tools to extract the timeseries in native EPI space between 100 cortical (Schaefer et al., 2018) and 16 subcortical brain nodes (Tian et al., 2020). Specifically, we co-registered the cleaned functional data to the T1-weighted structural image using FLIRT (Jenkinson and Smith, 2001). The T1-weighted image was then co-registered to the standard MNI space using FLIRT (12 DOF) and FNIRT (Andersson et al., 2007; Jenkinson and Smith, 2001). The resulting transformations were concatenated, inversed, and applied to warp the resting-state atlas from MNI space to the cleaned functional data in native space, preserving the labels with a nearest-neighbor interpolation method. Finally, we used custom-made Matlab scripts to extract the averaged time series using fslmaths and fslmeants.

### Probabilistic Tractography preprocessing

For each participant, a whole-brain structural connectivity matrix (SC) was computed using the atlas described above following the steps of previous studies (Muthuraman et al., 2016; Cao et al., 2013; Gong et al., 2009). The analysis was conducted using FMRIB’s Diffusion Toolbox (FDT) in FMRIB’s Software Library (FSL). First, the DICOM images were converted into Neuroimaging Informatics Technology Initiative (NIfTI) format using dcm2nii. The b0 image in native space was then co-registered to the T1-weighted image using FLIRT (Jenkinson and Smith, 2001). The co-registered T1 image was then co-registered to the standard space using FLIRT and FNIRT (Andersson et al., 2007; Jenkinson and Smith, 2001). The resulting transformation was inverted and applied to warp the atlas from the MNI space to the native MRI diffusion space using nearest-neighbor interpolation. Next, the diffusion-weighted images were processed using the FDT pipeline in FSL. Non-brain tissues were removed using the Brain Extraction Tool (BET) (Smith, 2002). The eddy current distortions and head motion were corrected using the eddy correct tool (Andersson and Sotiropoulos, 2016), and the gradient matrix was reoriented to correct for motion (Leemans and Jones, 2009). To model crossing fibers, BEDPOSTX was used, and the probability of multi-fibre orientations was computed to improve the sensitivity of non-dominant fibre populations (Behrens et al., 2007, 2003). Probabilistic Tractography was then performed for each participant in their native MRI diffusion space using the default settings of PROBTRACKX (Behrens et al., 2007, 2003). Finally, the connectivity probability *SC_np_* between nodes *n* and *p* were calculated as the proportion of sampled fibres in all voxels in node *n* that reach any voxel in node *p*. Since diffusion tensor imaging (DTI) does not capture fiber directionality, the *SC_np_* matrix was symmetrized by computing its transpose matrix *SC_pn_* and averaging both matrices.

### Leading Eigenvector Dynamics Analysis (LEiDA)

The first step was to define the empirical brain states of resting-state and meditation from a quantitative point of view, by using the LEiDA method (Cabral et al., 2017). As outlined in **Figure 1a**, in each parcellated brain area the blood oxygenation level-dependent (BOLD) time series were filtered (0.04-0.07 Hz) and Hilbert-transformed. For all participants in resting-state and meditation, at each repetition time (TR), a BOLD phase coherence matrix – dynamical functional connectivity dFC(*t*) – was calculated between each brain area pair *n* and *p*. This was done by calculating the cosine of the phase difference:

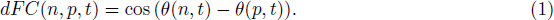

**Figure 1:**
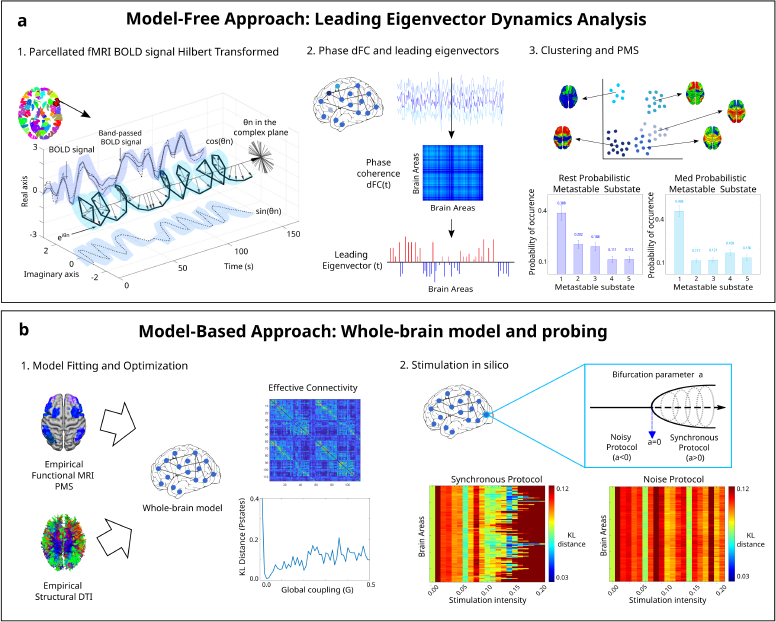
Methodology for model-free and model-based approaches. **a** Model-free framework: Leading Eigenvector Dynamic Analysis (LEiDA). In each parcellated brain area, the fMRI BOLD signal was band-passed filtered and Hilbert transformed to obtain the amplitude and phase information. For all participants in meditation and resting-state, the phase coherence matrix dFC(*t*) between brain areas was computed at each repetition time. The dimensionality of each matrix was reduced to its leading eigenvector *V*_1_(t) into positive (red color) and negative (blue color) values. Afterweards, K-means clustering was applied to the leading eigenvectors and the PMS computed for both meditation and resting-state. The optimal number of cluster centers was chosen according to the minimum value of *k* with the highest proportion of probabilities of occurrence of the metastable substates with significant difference between meditation and resting-state (*k* =5). **b** Model-based framework: whole-brain model. A Hopf whole-brain computational model was built for resting-state using the empirical frequency *w* calculated on the empirical fMRI signal, and fitting the modelled PMS space to the empirical PMS by looking at the global coupling *G* that minimized their KL distance. Effective connectivity was used to optimize the model by adjusting the DTI with a gradient descent approach until convergence. *In silico* stimulation was applied in each brain node separately by moving the bifurcation parameter *a* positively for a synchronization protocol, and negatively for a noise protocol. Optimal transition was detected for the bifurcation parameter that gave the minimum KL distance between the perturbed modelled PMS of resting-state, and the empirical PMS of meditation.

When a pair of nodes are temporarily aligned, the phase coherence is close to one since the difference between their Hilbert transformed signal angle is 0*^◦^* [cos(0*^◦^*)=1]. Contrarily, orthogonal BOLD signals have a phase coherence close to zero [cos(90*^◦^*)=0]. The resulting dFC(*t*) of each participant represented the interregional BOLD synchrony at each timepoint. The size of the matrix was N*×*N*×*T, where N is the number of brain areas (116) and T the total time points (220). A total of 880 (20 participants * 220 timepoints) matrices were calculated. A dominant connectivity pattern (dimension N*×*N) was obtained by reducing the dimensionality of the undirected and symmetric matrices into their leading eigenvectors *V*_1_(t) (dimension N*×*1) (Deco et al., 2019). The eigenvectors capture the dominant connectivity pattern at each time point *t*, representing the contribution of each brain area to the whole structure (Cabral et al., 2017). It is possible to reproduce the dominant connectivity pattern of dFC(*t*) by calculating the outer product of *V*_1_(t) and its transpose (V1.V1.*^T^*) (Lohmann et al., 2010). The leading eigenvectors dFC(*t*) for each TR and all participants from all states were then clustered with K-means clustering, varying *k* from 3 to 8. The resulting cloud centroids *V_c_*(t), which represent the dominant connectivity pattern in each cluster, were rendered onto a brain map using the Surf Ice software (https://www.nitrc.org/projects/surfice/).

Lastly, we computed the Probabilistic Metastable Substate Space (PMS), which expresses the probability of being in each metastable substate from the substate repertoire. Specifically, the ratio of number of epochs (TRs) in a cluster divided by the total number of epochs in all clusters was computed separately for resting-state and meditation.

### Whole-Brain Computational Model

A whole-brain computational model was built for resting-state using the normal form of supercritical Hopf bifurcation. This Landau-Stuart oscillator, referred to as Hopf, has been used to study transitions from noisy to oscillatory regimes (Deco et al., 2015) (**Figure 1b**). Emergent brain dynamics were simulated by linking the structural and functional connectivity using effective connectivity (Deco et al., 2015). For each of the 100 cortical and 16 subcortical brain areas (Tian et al., 2020; Schaefer et al., 2018), BOLD activity was emulated and the working point was fitted and optimized with specific parameters of the model (Deco et al., 2019).

The following pair of coupled equations can be used to represent an uncoupled node *n* in Cartesian coordinates:

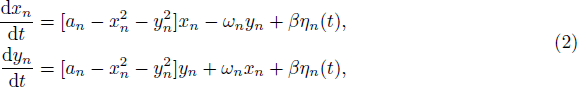

where *x_n_* represents the BOLD signal, *η_n_*(*t*) corresponds to the additive Gaussian noise, *β* = 0.01 to the standard deviation of the additive Gaussian noise, and *f_n_* = *ω_n_/*2*π* is the frequency of the system. This frequency was estimated by averaging the empirically filtered BOLD signals in the 0.04-to 0.07-Hz band for each brain node *n*=1,…, 116 (Deco et al., 2019). The parameter a=0 indicates the bifurcation point between noise and oscillations. The noise regime is found in *a<*0, whereas synchronous oscillations in *a>*0 at a firing frequency of *w/2π* (Deco et al., 2017a). A value of *a_n_*=-0.02 for each brain node *n* was selected based on previous findings (Deco et al., 2017b).

Furthermore, a global coupling weight *G* and an additive coupling term *C_np_* were included. The whole-brain dynamics at each node *n* were defined as:

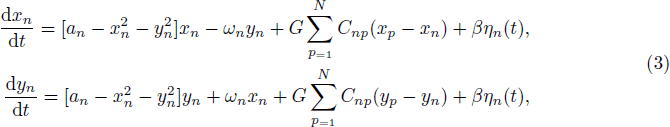

where the global coupling weight *G* represents the strength between all nodes (i.e., axonal conductivity) and scales all of the connections equally. The additive coupling term *C_np_* is built upon the SC and represents the input to node *n* from each of the rest of the nodes *p* (i.e., myelination density). It is a weighted matrix since it assumes different densities for each node.

#### Model Fitting: Comparing empirical and simulated probability metastable space states

The global coupling weight *G* was ranged from 0 to 0.5 in steps of 0.01 and the model run 200 iterations for each value of *G*. In order to obtain the simulated PMS space, LEiDA was computed on the Hilbert transformed simulated signal based on the centroids already defined in the empirical analysis. Maximal model fit was obtained for the value of *G* that resulted in a maximal similarity between the modelled PMS at resting-state and the empirical PMS at resting-state, corresponding to their lowest Kullback-Leibler (KL) distance (Deco et al., 2019). This metric is given by:

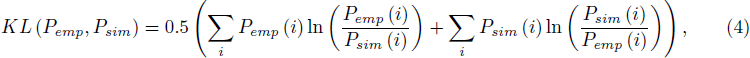

where for each brain metastable substate *i*, *P_emp_*(*i*) are the empirical probabilities and *P_sim_*(*i*) the simulated probabilities.

#### Model Optimization: Method for updating Effective Connectivity

For each value of *G*, the SC was updated to address potential connections that were missing. The initial value of *C* corresponded to the average of the empirical DTI structural connectivity of all participants normalized to 0.2 (Deco et al., 2019, 2017b, 2015). The distance between the grand average phase coherence matrices of the model 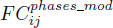 and the empirical matrices 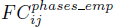 was calculated iteratively and the SC was transformed to effective connectivity (EC). A gradient descent approach adjusted each structural connection between each pair of nodes *i* and *j* :

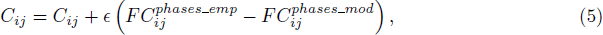

in which *ɛ* = 0.01, and the grand average phase coherence matrices is given by:

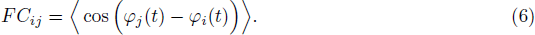

The Hilbert transform BOLD signal phase of nodes *j* and *i* at time *t* is expressed by *φ*(*t*) and the brackets indicate the average across time. Each time, the empirical and simulated values were compared until the difference was less than 0.001(Deco et al., 2019).

### Unilateral Perturbation of the Whole-Brain Model

Transitions from resting-state towards the target meditative state were evaluated as schematized in **Figure 1b**. The whole-brain model for the resting-state condition was perturbed unilaterally by moving individually and separately the local bifurcation parameter *a* of each of the 116 brain areas. A measure of the fit to the target state (empirical meditation) was evaluated by computing the KL distance between the perturbed modelled resting-state PMS and the target empirical meditation PMS. Areas with the lowest KL distance were predicted to promote transitions after simulation. There were two different protocols studied, synchronization and noise. As for the synchronization protocol, the bifurcation parameter was shifted from 0 to 0.2 in steps of 0.01; as for the noise protocol, it was shifted from 0 to -0.2 in steps of -0.01. Stimulation intensities are represented by the absolute value of each step (Deco et al., 2017b). In order to minimize random effects from the Gaussian noise of the model, each simulation was repeated 3 times (Deco et al., 2019).

### Statistical Analysis

The statistical analysis were performed using MATLAB R2022a from MathWorks (Natick, MA, USA). To test the results of the PMS obtained by LEiDA, Wilcoxon tests with 1000 iterations were used to examine the probability of occurrence of all explored clustering conditions (*k* from 3 to 8). Permutations were compared using the Wilcoxon test with a significance threshold of 0.05. Multiple comparisons were corrected with the False Discovery Rate (FDR) method (Hochberg and Benjamini, 1990) when testing the differences between resting-state and meditation for all metastable substates in each value of *k*. The reported P-values remain significant after FDR correction.

## Results

### LEiDA

The most accurate description of the empirical data across participants corresponded to a value of cluster centers *k* =5. This amount of clusters was the minimum number of *k* that provided a maximal proportion of statistically significant differences between resting-state and meditation, surviving correction by multiple comparisons. We illustrate the probability of occurrence of the PMS of resting-state and meditation in **Figure 2a**. In addition, we rendered cluster centroid eigenvectors onto the brain maps in **Figure 2b**. The sign of eigenvector elements are positive or negative and indicated by red and blue colors, respectively. The signs represent the direction that the BOLD signals from different elements follow in relation to their projection onto the leading eigenvector. Hence, classifying elements into communities in which color strength reflects how strongly each area is associated with its community (Alonso Martinez et al., 2020; Cabral et al., 2017). In the first metastable substate, all eigenvector elements had the same sign. On the other hand, substate 2 showed a functional network dominated mainly by the somatomotor and salience networks, and the putamen. Additionally, substate 3 displayed a community comprised mostly of the DMN and a few areas from the limbic system, whereas substate 4 showed coordination primarily between the control network and the DMN. Lastly, substate 5 had a network principally led by the visual system and some areas of the somatomotor and dorsal attention networks. The probability of occurrence of meditation compared to resting-state was significantly higher in substate 4 [0.169 ± 0.021 vs. 0.111 ± 0.019, *P* =0.019], and lower in substate 2 [0.117 ± 0.015 vs. 0.202 ± 0.023, *P* =0.008] and substate 3 [0.121 ± 0.015 vs. 0.188 ± 0.023, *P* =0.025].

**Figure 2:**
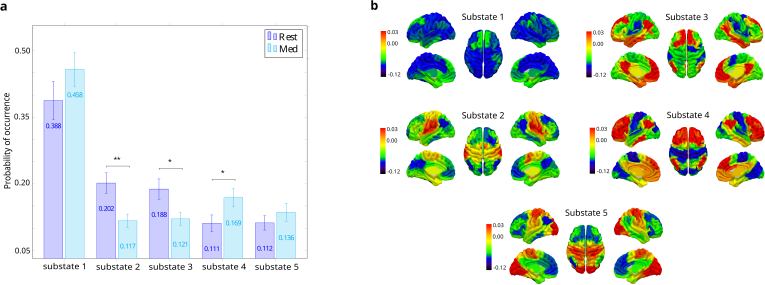
**a** Empirical Probabilistic Metastable Substate (PMS) Space. Mean probability of occurrence of resting-state and meditation in each metastable substate. Differences were computed with a 95% confidence interval and significance is represented with asterisks (* p *<* 0.05, ** p *<* 0.01 and *** p *<* 0.001). Substates 1, 3 and 4 had a higher probability of occurrence during meditation compared to resting-state. Substates 2 and 3 presented the opposite behavior. **b** Cluster centroids *Vc*(t) rendered onto brain maps, representing leading eigenvectors. Substate 1 had all eigenvector elements with the same sign. Substate 2 was characterized with an interaction mostly between the somatomotor network, salience network, and the putamen. Substate 3 presented a functional network dominated by the DMN and areas from the limbic system. Substate 4 had community formed mainly by the control network and the DMN. Substate 5 showed positive values mostly in areas from the visual system, and also the somatomotor network and dorsal attention network.

### Whole-brain computational model for resting-state

We fitted and optimized a whole-brain model for resting-state in order to find the parameters that maximally approximated its empirical PMS, corresponding to the minimal KL distance between the simulated and the empirical PMS. The best fit was found at a global coupling of *G* =0.03 with a KL distance of 0.008 (**Figure 3a**). The similar probabilities of occurrence for all of the five substates are shown in **Figure 3b**.

**Figure 3:**
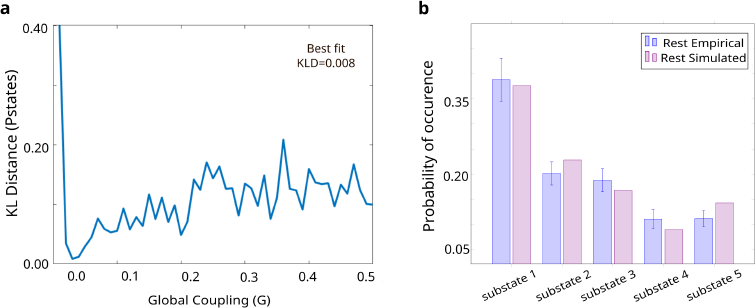
Model-based results: Whole-brain model. **a** Optimal fit and optimization for resting-state model at a global coupling weight *G* of 0.03. **b** Empirical and simulated PMS for resting-state in *G*=0.03.

### *In silico* stimulations to force transitions from resting-state to meditation

In the last step of the analysis, we systematically perturbed the model of resting-state and compared the resulting perturbed PMS with the empirical PMS of meditation. For the synchronous protocol, we shifted the bifurcation parameter *a* independently in each brain area, with positive values. Contrarily, for the noise protocol, we used negative values. We identified the optimal perturbation as the one resulting in the smallest KL distance between the perturbed modelled source PMS (resting-state) and the target empirical PMS (meditation).

Results for perturbing the whole-brain model of resting-state in the synchronization and noise protocols at different stimulation intensities are illustrated in **Figures 4a** and **4b**, respectively. The color-scale indicates the KL distances between the empirical and the perturbed PMS after stimulating each brain area separately. The noise protocol was not successful in transitioning from resting-state to meditation. In contrast, in the synchronization protocol, KL distances decreased with increasing positive bifurcation point values *a*. The lowest KL distances for the maximal number of brain areas were given at a perturbation value of *a*=0.14. At this stimulation intensity, the most sensitive areas belonged mainly to somatomotor and dorsal attention networks, and fewer areas of the salience, visual and default-mode networks. For values of *a>*0.14, the system drifted again away from the target meditative state.

**Figure 4:**
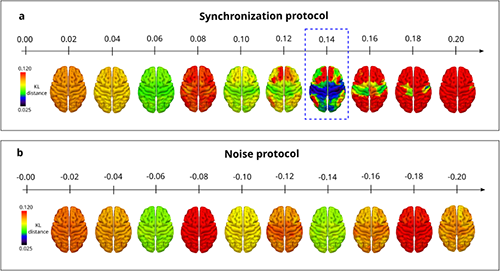
Model-based results: *In silico* stimulation. We stimulated one brain area at a time to study whole-brain effects and evaluate a possible transition from resting-state to meditation. The local bifurcation parameter *a* was shifted independently for each node. The x-axis represents stimulation intensities and the color scale shows the KL distance between the perturbed PMS and the target PMS. **a** In the synchronization protocol, the system moved closer to the meditation state for positive values of *a* with an optimal transition in *a*=0.14 for most of the areas. The best fit was obtained when perturbing the RH Som 6. For values higher than *a>*0.14 the system drifted away. **b** In the noise protocol, the KL distance between the perturbed modelled PMS of resting-state and the empirical PMS of meditation, never decreased for increasing negative values of *a*.

The best transition for each brain area was obtained at the stimulation intensity which provided the minimum KL distance between the perturbed PMS and the target PMS, illustrated in **Figure 5a**. The minimum KL distance for the optimal perturbation of each brain area is rendered into the brain maps of **Figure 5b**. The top 10 areas to perturb, at their particular optimal stimulation intensity in the synchronization protocol, belonged to the somatomotor network and dorsal attentional network. We achieved the best fit when perturbing the right somatomotor cortex (RH Som 6 from the Schaeffer 100 parcellation (Schaefer et al., 2018)) at an intensity of *a*=0.14. Consequently, the perturbed and target PMS were very similar, as reflected in **Figure 5c**.

**Figure 5:**
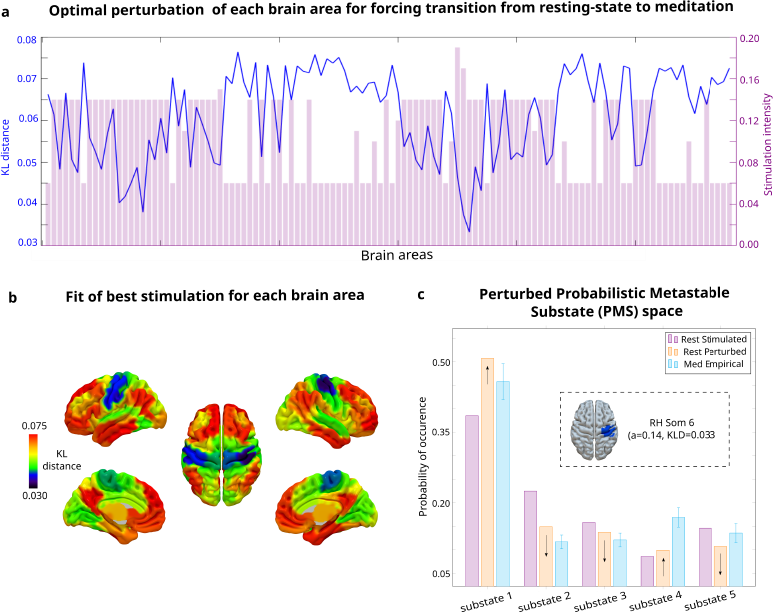
Optimal perturbational results. **a** Best fit for forcing transition from resting-state to meditation. Optimal perturbation for each brain (x-axis) area in the synchronization protocol, found at the stimulation intensity (right y-axis) which gives the minimum KL between the modelled PMS of resting-state after perturbation, and the empirical PMS of meditation (left y-axis). Optimal fit was found when stimulating the RH Som 6 with an intensity of *a*=0.14. **b** Fit of optimal stimulation for each brain area in the synchronization protocol. Colorcode represents the KL distance between the perturbed modelled PMS of resting-state and the empirical PMS of meditation. **c** PMS for simulated resting-state, perturbed resting-state and empirical meditation. When the resting-state model was perturbed at the RH Som 6 with an intensity of *a*=0.14, the probabilities of occurrence of substates 1 and 4 increased, whereas for substate 2, 3 and 5 decreased.

## Discussion

We successfully studied brain dynamics using model-free and model-based approaches (Deco et al., 2017b) to find causal mechanisms explanations for meditation in a group of expert meditators. LEiDA allowed us to characterize brain signatures of meditation and resting-state by defining their PMS, with an associated probability of occurrence of being in distinct metastable substate (i.e., dynamic pattern) (Cabral et al., 2017). Interestingly, we found brain dynamics differences between resting-state and meditation in areas that overlap with the triple-network model (Menon, 2011). Then, we fitted a Hopf model to the empirical PMS of resting-state. Consequently, this allowed us to force a transition from the PMS of the resting-state model to the PMS of the target meditation using off-line *in silico* unilateral perturbations. By varying stimulation intensities and protocols, we were able to determine the mechanistic global effects of exhaustive local perturbations. This way, we revealed how local changes in resting-state can alter whole-brain dynamics and evaluated the optimal transition towards a meditative state. Using a synchronization protocol, we could transition from the resting state to the meditative state. The most sensitive brain regions were located in somatomotor and dorsal attention networks.

In the model-free approach, we used LEiDA to reveal signatures of interacting brain networks for meditation and resting-state (**Figure 2**). During meditation, compared to the resting state, we found a significantly lower probability of occurrence in substates 2 and 3, and a significantly higher probability in substate 4. Firstly, we found that substate 2 exhibited a functional network predominated by the somatomotor network, the salience network and the putamen. The somatomotor network is responsible for processing bodily sensations and executing reactions (ten Donkelaar et al., 2020), and it has been shown that its activity decreases in meditation (Jerath et al., 2012). Additionally, the salience network monitors reaction to external information, switching between internal and external processing to external stimuli, and processing of pain, emotion, reward and motivation (Goulden et al., 2014; Elton and Gao, 2014). A reduction in functional connectivity within the salience network has been associated with experiential acceptance and present-moment awareness (Sezer et al., 2022). Interestingly, Guidotti et al. (2023) used a trained classifier revealing that this network was able to discriminate meditation style in an expert group, highlighting its crucial role in specific meditation techniques. Lastly, the putamen, involved in motor control and learning (Viñas-Guasch and Wu, 2017), has been related to the attentional disengagement from irrelevant information during meditation in expert meditators Sperduti et al. (2011). With regards to substate 3, we found it displayed a community led majorly by the DMN and some areas of the limbic system. Our result aligns with previous studies reporting a reduction in DMN activity during meditation Ganesan et al. (2023); Garrison et al. (2015), which could be associated with the phenomenological well-being of meditation (Brandmeyer and Delorme, 2021; Killingsworth and Gilbert, 2010). On the other hand, we revealed that substate 4 presented a coordination mainly between the central executive network and the DMN. The central executive network is associated with executive functions (i.e., attention, cognitive control and working memory) (Niendam et al., 2012), whereas the DMN is involved in internal thought, self-related processing and mind-wandering (Alves et al., 2019; Mason et al., 2007). Their interaction, therefore, is necessary for balancing self-referential and external stimuli processing (Chen et al., 2013). Bauer et al. (2019) reported increased connectivity between areas of the DMN and frontoparietal network in meditators during a practice suggesting a top-down regulation from the CEN to the DMN in order to suppress distraction and direct the focus of attention. Furthermore, Kajimura et al. (2020) reported the integration of the frontoparietal network into the DMN in a single naive participant after 65 days of meditation, explained by the inhibition of the DMN during a focus task which increases concentration and emotion regulation.

Interestingly, we found an overlap between some leading networks in the metastable substates revealed in LEiDA and the networks from the triple-network model: the DMN, the salience and the central executive network (Menon, 2011). This model identifies the salience and central executive network as task-positive networks and the DMN as a mind-wandering network. A modulator role is given to the salience network, responsible for evaluating sensory inputs and switching the activations of the central executive network and the DMN when in an attentional goal-oriented state or at rest, respectively. The relevance of self-referential and attentional network in meditation studies has been highlighted by several research groups (Bremer et al., 2022; Vishnubhotla et al., 2021; Lim et al., 2018; Gotink et al., 2016; Berkovich-Ohana et al., 2012). An interesting study conducted by Hasenkamp et al. (2012) evaluated breath-focused meditation in experienced meditators, which had to press a button when they noticed focus was lost (i.e., mind-wandering). This way, researchers identified a cognitive cycle of four stages and fMRI activation patterns: 1) loss of focused attention was associated with DMN activation, 2) awareness of mind-wandering coexisted with salience network activation, 3) shift of attention from mind-wandering and redirection of attention back to the breath happened when the central executive network activity increased, and lastly, 4) a sustained attention to breath coexisted with activity in the central executive network (in particular the dorsolateral prefrontal cortex). The relevance of this DMN-salience-central executive network interplay comes from the fact that it is increasingly explored in brain disorders (e.g., schizophrenia, attention deficit hyperactivity disorder, depression) (Menon, 2011) and developmental studies in children (Denervaud et al., 2020; Ryali et al., 2016). Therefore, regular practice of meditation could help re-balance the interaction between these networks (Bremer et al., 2022) as well as develop cognitive skills related to attention maintenance and disengagement from distraction (Hasenkamp and Barsalou, 2012; Hasenkamp et al., 2012).

For the model-based approach, we first built a Hopf whole-brain computational model of resting-state, which allowed us to systematically evaluate *in silico* stimulations to force a transition to a meditative state (Deco et al., 2017b). We obtained the resting-state model by optimizing empirical functional and structural data and determining a global coupling value of *G* =0.03, corresponding to the lowest KL distance between the empirical PMS and the modeled PMS (**Figure 3**). We then explored local perturbations region by region in order to study their dynamical causal whole-brain effects as well as the possibility of shifting the system from resting-state to a meditative state. We could transition successfully from resting-state to meditation with a synchronous regime, led by oscillations, and not in a noisy protocol (**Figure 4**). Furthermore, we approached the meditative target state with increasing positive stimulation intensities, reaching an optimal transition at a bifurcation point of *a*=0.14, and then drifted away. Our results show relevant areas most sensitive to perturb were found in the somatomotor and dorsal attentional networks (**Figure 5**). The importance of the former comes from the fact that during meditation body awareness processing is altered (Tang et al., 2015) and attention has to be oriented to the target stimulus – interoceptive sensation (i.e., breathing) – and disengage from non-target stimuli – distractors (Ganesan et al., 2022; Kabat-Zinn, 2003). In particular, our results showed that the most optimal area for perturbation was in right somatomotor cortex (RH Som 6 from the Schaeffer 100 parcellation (Schaefer et al., 2018)) at a stimulation intensity of *a*=0.14. Overall, our model-based results provide novel insights into how local perturbations influence global brain dynamics. We could explore the most sensitive brain areas in terms of their response to perturbations as a model-based biomarker of the underlying mechanisms of meditation.

This study has some limitations. Firstly, we analyzed a relatively small sample size dataset. Therefore, validating our results using a larger dataset would be beneficial (Fox et al., 2016). In addition, our analysis focuses on a specific meditation technique which prevents the generalization of our findings to all styles of meditation. Given the heterogeneity of the practice, we cannot oversimplify our results to one signature of brain dynamics in meditation (Young et al., 2021; Fox and Cahn, 2019). Moreover, our study solely examines state differences between resting-state and meditation, specifically in expert meditators. In order to study the effects of meditation practice on brain structure and function over time, longitudinal studies are necessary. As a remark, we want to emphasize the importance of not oversimplifying the Eastern tradition of meditating to a single motivation around attention and emotion regulation (Garcia-Campayo et al., 2021). Additionally, if used as a therapeutic intervention, it is essential to recognize that meditation is not a cure-all solution (Wright, 2013).

In this work, we revealed the underlying brain mechanisms in a group of expert meditators during focused attention on breathing. Using the LEiDA model-free approach, we could quantitatively define the brain dynamics during meditation and resting-state and found differences in areas mainly overlapping the triple-network model (Deco et al., 2019; Cabral et al., 2017). In addition, the model-based approach allowed us to measure the whole-brain reactivity to localized perturbation at resting-state and study transitions towards meditation. Successful transitions were found in a synchronization protocol and the most sensitive areas were located in the somatomotor and dorsal attention networks. Given the paucity of research in meditation with neuroscience tools, our results shed light on the field of contemplative neuroscience for promoting meditation practices in health and a potential therapy in disease (Ludwig and Kabat-Zinn, 2008; Grossman et al., 2004). Our findings could open future investigations in meditation research and other non-ordinary states of consciousness.

## Data availability

Raw data for the current study may be obtained from the corresponding author upon reasonable request.

